# Single-nucleus analysis reveals oxidative stress in Down syndrome basal forebrain neurons at birth

**DOI:** 10.1101/2025.02.05.636750

**Authors:** Nicole R. West, Kalpana Hanthanan Arachchilage, Sara Knaack, Shawn MacGregor, Masoumeh Hosseini, Ryan D. Risgaard, Pubudu Kumarage, Jose L. Martinez, Su-Chun Zhang, Daifeng Wang, Andre M.M. Sousa, Anita Bhattacharyya

## Abstract

**INTRODUCTION:** Basal forebrain cholinergic neurons (BFCNs) are integral to learning, attention, and memory, and are prone to degeneration in Down syndrome (DS), Alzheimer’s disease, and other neurodegenerative diseases. However, the mechanisms that lead to the degeneration of these neurons are not known.

**METHODS:** Single-nucleus gene expression and ATAC sequencing were performed on postmortem human basal forebrain from unaffected control and DS tissue samples at 0-2 years of age (n=4 each).

**RESULTS:** Sequencing analysis of postmortem human basal forebrain identifies gene expression differences in DS early in life. Genes encoding proteins associated with energy metabolism pathways, specifically oxidative phosphorylation and glycolysis, and genes encoding antioxidant enzymes are upregulated in DS BFCNs.

**DISCUSSION:** Multiomic analyses reveal that energy metabolism may be disrupted in DS BFCNs by birth. Increased oxidative phosphorylation and the accumulation of reactive oxygen species byproducts may be early contributors to DS BFCN neurodegeneration.

## 1. Background

Cholinergic projection neurons of the basal forebrain (BFCNs), are the primary cholinergic input to the cortex, hippocampus, and amygdala, regulating cognitive functions including learning, attention, and memory(1). The cholinergic hypothesis, proposed nearly 50 years ago, posits that the dysfunction or loss of cholinergic neurons is an early driver of cognitive decline associated with age and Alzheimer’s disease (AD)(2–4). BFCNs are some of the first neurons to degenerate in the progression of AD(5–7). Tau tangles accumulate in the basal forebrain in AD before the entorhinal cortex(5, 6, 8). Studies suggest that BFCNs seed the cortex with pathology through the trans-synaptic spread of misfolded Tau(6, 7). Consequently, BFCNs have been a target of therapeutics to reduce degeneration, slow the spread of AD pathology, and ultimately to slow cognitive decline in AD(9, 10). Although the link between BFCN degeneration and memory decline is well understood, early molecular events that occur in BFCNs that increase susceptibility to degeneration later are unknown.

BFCN dysfunction and degeneration also occur in Down syndrome (DS, trisomy 21, T21)(11–15) and other neurodegenerative diseases, including Parkinson’s disease (PD)(12, 16–20) and Dementia with Lewy bodies (DLB)(21–24). Nearly all individuals with DS develop AD (DS-AD), making DS the leading genetic cause of AD(25). The progression of DS-AD pathology and the onset of dementia occurs in a consistent and predictable manner(26, 27), enabling examination of successive stages of disease progression in DS. BFCN degeneration in the anteromedial and posterior basal forebrain in individuals with DS begins about 30 years prior to the median onset of prodromal AD(14). Post-mortem studies validate fewer neurons are present in the basal forebrain in DS relative to unaffected controls(11). Further, DS is a neurodevelopmental disorder, and so we hypothesized that early molecular changes in BFCNs increase their susceptibility to degeneration in DS. Identifying these molecular changes may provide insight into drivers of BFCN degeneration in the progression of DS-AD.

We performed unbiased single-nucleus gene expression and ATAC multiomic analysis of early human postnatal DS and unaffected control basal forebrain (BF) to identify molecular changes in DS that precede BFCN dysfunction or degeneration(14, 28–30). We identified cell type-specific differential gene expression across all eight cell types in the DS basal forebrain. The results suggest that energy metabolism, specifically the upregulation of glycolysis and oxidative phosphorylation (OXPHOS) genes, is dysregulated in DS BFCNs. Consequently, DS BFCNs have increased gene expression of antioxidant enzymes, possibly to regulate the reactive oxygen species (ROS) that accumulate as a byproduct of OXPHOS. The increased ROS over a sustained time may increase the vulnerability of DS BFCNs, causing their early degeneration in DS. These results identify potential novel targets and a timeframe for therapeutic intervention to delay BFCN dysfunction and mitigate disease progression.

## 2. Methods

### 2.1 Tissue Samples

Human basal forebrain post-mortem samples from Down syndrome and unaffected control individuals were obtained from the University of Maryland Brain and Tissue Bank, as part of the National Institutes of Health NeuroBioBank. Acquisition of the de-identified samples was approved by the Health Sciences Institutional Review Board at the University of Wisconsin-Madison (Protocol #2016-0979) and certified exempt from IRB oversight. Sample information is provided in **Figure 1A** and **Supplement Table 1**.

**Figure 1.**
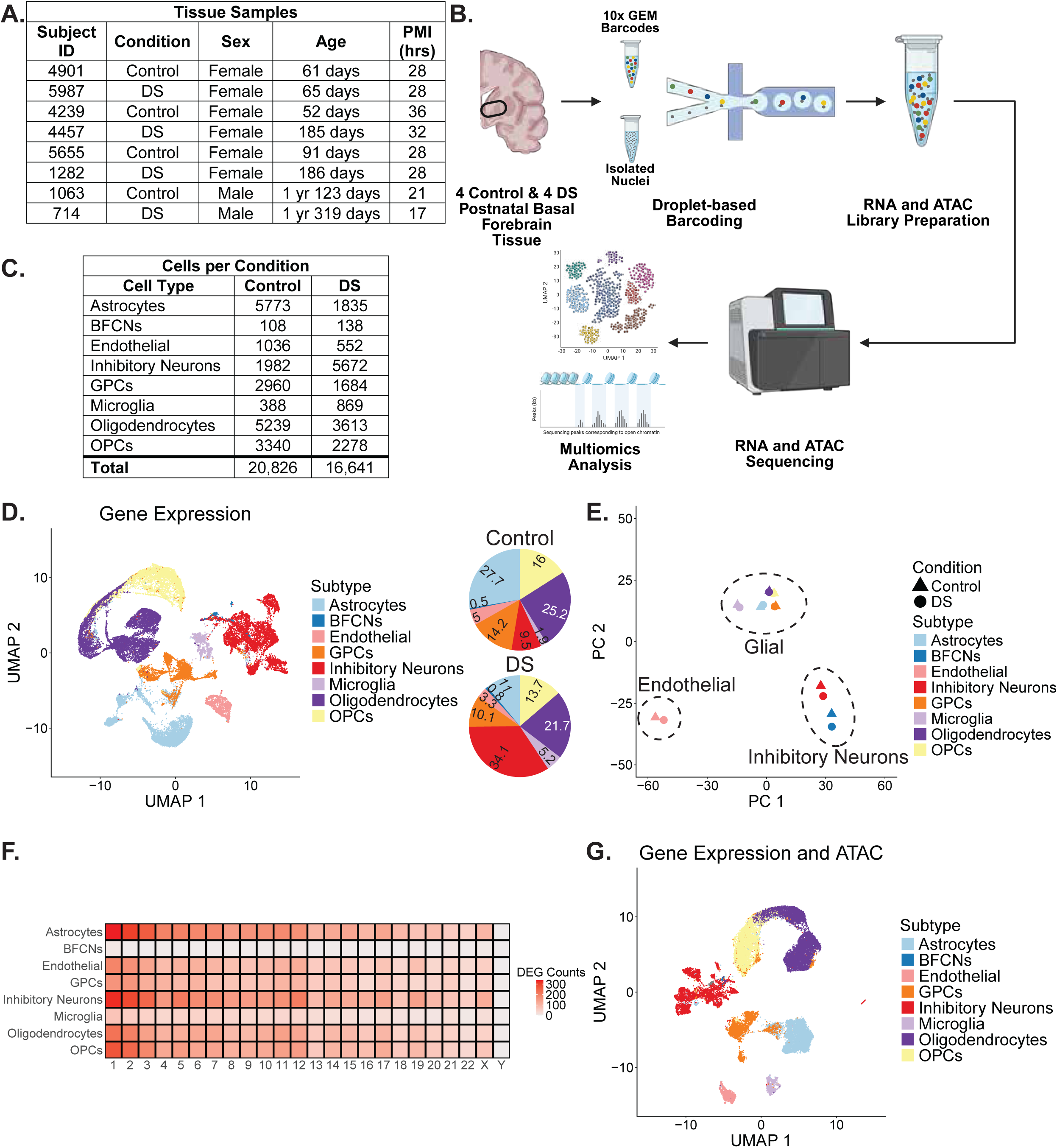
A) BF samples from four control and four DS donors matched for age, sex, and PMI were used in this study. B) Schematic of BF snMultiomic analysis (Created with BioRender.com). C) Cell types and the numbers identified in control and DS samples. D) Gene expression UMAP of cell clusters by subtype. Cell subtypes identified in the human basal forebrain were astrocytes, BFCNs, endothelial cells, GPCs, inhibitory neurons, microglia, oligodendrocytes, and OPCs. Percentages of cell types in control (N=20,826) and DS (N=16,641). There are fewer astrocytes and more inhibitory neurons and microglia in the DS basal forebrain. E) PCA analysis of cell type and genotype. Cell types from control and DS are largely the same, with control and DS clustering together on the plot. Cell subtypes cluster by subclass. F) DEGs per chromosome by cell type. DEGs are distributed across the genome in all cell types. G) Gene expression and ATAC integrated UMAP of cell clusters by subtype.

### 2.2 Nuclei Isolation

Frozen post-mortem human basal forebrain (BF) tissue sections were pulverized into a powder in liquid nitrogen over dry ice, using a mortar and pestle (Fisherscientific #FB961A and #FB961K). 25-35 mg of pulverized BF tissue was used for nuclei isolation. All buffers were prepared fresh and maintained on ice. 3 mL of ice-cold buffer B (Iodixanol buffer) [50% (v/v) Optiprep (Iodixanol) solution (Sigma # D1556); 25 mM KCl (Sigma #60142); 5 mM MgCl2 (Sigma #M1028); 20 mM Tris-HCl (pH 7.5) (Invitrogen #15567-27); 1% cOmplete™, Mini, EDTA-free Protease Inhibitor Cocktail (Roche #11836170001); 1% BSA (GeminiBio #700-100p); RNase inhibitor (80U/mL) (Roche #03335402001); 1 mM DTT (Sigma #43186)] was transferred to a 15 mL tube.

1 mL buffer A (lysis buffer) [250 mM Sucrose (Sigma #S0389); 25 mM KCl2 (Sigma #21115); 5 mM MgCl2 (Sigma #63052); 20 mM Tris-HCl (pH 7.5) (Invitrogen #15567-027); 1% cOmplete™, Mini, EDTA-free Protease Inhibitor Cocktail (Roche #11836170001); RNase inhibitor (40U/mL) (Roche #03335402001); 1 mM DTT (Sigma #43186); 0.1 % (v/v) IGEPAL CA-630 (Sigma #I8896)] was added to the pulverized BF tissue in a tube. 1 mL of ice-cold lysis buffer was added to the Dounce tissue grinder (15 mL volume, Wheaton #357544; autoclaved, RNase free, ice-cold). 1 mL of lysis buffer was added to the pulverized tissue tube to rinse and collect all tissue. The suspension was transferred to Dounce tissue grinder and homogenized with loose and tight pestles, 30 cycles each, with constant pressure and without the introduction of air. The solution was transferred to the 15 mL tube containing buffer B and mixed by inverting the tube 10 times. The homogenate was filtered through a 40-um cell strainer (Falcon #352340) which was pre-washed with lysis buffer. Samples were centrifuged at 1000 g for 30 min at 4°C (Eppendorf #5910 Ri). Following centrifugation, the supernatant was carefully and completely removed and the pellet was resuspended in 1 mL of wash buffer [10 mM NaCl (Sigma #60142); 3 mM MgCl2 (Sigma #M1028); 10 mM Tris-HCl (pH 7.5) (Invitrogen #15567-027); 1% BSA (GeminiBio #700-100p); RNase inhibitor (1000U/mL) (Roche #03335402001); 1mM DTT (Sigma #43186)]. The homogenate was filtered through a 40-um cell strainer (Falcon #352340) pre-washed with wash buffer to eliminate large clumps and cell debris. Samples were then centrifuged at 500 g for 5 min at 4°C.

Supernatants were carefully and completely removed. Pellets were gently dissolved by adding 200 mL and 300 mL of buffer C [10 mM NaCl (Sigma #60142); 3 mM MgCl2 (Sigma #M1028); 10 mM Tris-HCl (pH 7.5) (Invitrogen #15567-027); 0.01% Tween-20 (Bio-Rad #1662404); 0.001% Digitonin (Thermo Fisher #BN2006); 0.01% (v/v) IGEPAL CA630 (Sigma#I8896); 1% cOmplete™, Mini, EDTA-free Protease Inhibitor Cocktail (Roche#11836170001); 1% BSA (GeminiBio #700-100p); RNase inhibitor (1000U/mL) (Roche #03335402001); 1mM DTT (Sigma #43186)].The solution was incubated on ice for 5 minutes. After incubation 500 mL of buffer D [10 mM NaCl (Sigma #60142); 3 mM MgCl2 (Sigma #M1028); 10 mM Tris-HCl (pH 7.5) (Invitrogen #15567-027); 0.1% Tween-20 (Bio-Rad #1662404); 1% BSA (GeminiBio #700-100p); RNase inhibitor (1000U/mL) (Roche #03335402001); 1mM DTT (Sigma #43186)) was added to the solution. After resuspension, nuclei quality was assessed at 40X magnification and were manually counted using a hemocytometer. The sample was centrifuged at 500 g for 5 min at 4°C. The pellet was resuspended in buffer E [1X nuclei buffer (10x Genomics #2000207); RNase inhibitor (1000U/mL) (Roche #03335402001); 1mM DTT (Sigma #43186)] at a final concentration of 5 million nuclei/mL.

### 2.3 snMultiomic Library Generation and Sequencing

For each sample, 10,000 nuclei were targeted. Nuclei suspension was first incubated in a transposition mix. Thereafter, along with the oligo-coated gel beads and partitioning oil (10x Genomics #PN-2000190), the single nuclei master mixture containing tagmented single nuclei suspension was transferred onto a Next GEM Chip J (10x Genomic #PN-2000264), and the chip was loaded to the Chromium Controller for GEM generation. After post GEM incubation clean up, preamplification of samples was performed and ATAC libraries were generated utilizing the Single Index Kit N Set A (10x Genomics #PN-1000212). The snRNA-seq libraries were generated using the Library Construction Kit (10x Genomics #PN-1000190) and Dual Index Kit TT Set A (10x Genomics #PN-1000215), following the manufacturer’s recommended protocol. At each step, the concentration and quality of cDNA, ATAC library and GEX libraries were assessed by 4200 TapeStation (Agilent). Sequencing was carried out with Illumina NovaSeq X Plus for a targeted depth of 50,000 raw reads per nucleus.

### 2.4 snMultiomic Alignment

The 10x multiomic data was processed according to BICAN default methods. The snRNA-seq data were aligned with STAR v2.7.11a(31) and aggregated into count matrices with STARSolo. The alignments were performed locally using defaults from the WARP Multiome (v5.9.0) pipeline(32). Initial quality control checks included assessment of the percentages of uniquely- and multiply-mapped reads, and statistics of corrected barcodes and UMI. The snRNA-seq data alignment was done utilizing the GeneFull_Ex50pAS argument for STAR to ensure inclusion of alignments that overlap exonic ends within genic regions, in consideration of nuclear RNA processing biology.

The snATAC-seq data was likewise processed according to the WARP Multiome (v5.9.0) pipeline. This includes ibarcode correction with the fastqProcessing processing tool from WarpTools (32), and subsequent trimming with Cutadapt (v4.4)(33). Alignment was performed with BWA-MEM2(34). Finally, generation of fragment files and initial QC was performed with SnapATAC (v2.3.1)(35). This was facilitated with an updated version of the snATAC component of the WARP Multiome (v5.9.0) pipeline to facilitate this workflow on Amazon AWS EC2 instances. Specifically, docker images for Cutadapt (v4.4) and for the BWA alignment were prepared to leverage AVX2 processing for improved performance. In this stage of the analysis all samples for either unaffected control or DS donors were treated identically.

### 2.5 snRNA-seq Processing and QC

Data processing and downstream analysis were conducted in R (4.4.0). Packages and versions are listed in **Supplement Table 2**. Since DS samples carry an extra chromosome, we modified the standard preprocessing pipeline to incorporate genotype-specific differences in the transcript abundance. For instance, it is standard practice to impose upper and lower thresholds on the number of UMIs (nCount_RNA). The upper threshold is primarily used to remove potential homotypic doublets. Applying a single upper UMI threshold would have either led to a loss of many high-quality DS cells or the retention of potential doublets in control samples. To address this, we set separate upper UMI thresholds for control and DS samples based on their respective distributions (**Supplement 1A**). Identifying highly variable genes separately per genotype enhances the transcriptomic signal related to batch effects and cell types, ultimately improving clustering accuracy.

The snRNA-seq datasets were preprocessed and analyzed using the Seurat (v5.1.0)(36, 37) R package. mRNA contamination caused by cell-free ambient RNA in the gene expression data was corrected using the SoupX (v1.6.2)(38) package. Low-quality nuclei were then identified and removed based on stringent quality control thresholds: fewer than 200 expressed genes, ribosomal gene content exceeding 40%, mitochondrial gene content exceeding 5%, and a UMI count lower than 800 or higher than 𝑄_3_ + 3(𝑄_3_ − 𝑄_1_), where 𝑄_1_ and 𝑄_3_ are the lower and upper quartiles(39, 40) (**Supplement 1A-B**). The upper UMI thresholds used were 29,284 in control and 34,160 in DS (**Supplement 1A**). After quality control, the datasets were subjected to doublet removal. Given the uncertainties inherent in doublet detection methods, an ensemble approach was employed, incorporating three techniques: DoubletFinder (v2.0.4)(41), Scrublet(42), and scDBIFinder v(1.18.0)(43). A cell identified as a doublet by at least two of the three methods was classified as a doublet and removed from the dataset.

SoupX-corrected UMI counts were log-normalized using Seurat. The top 3,000 highly variable genes were identified using the default variance-stabilizing process. Gene expression data for these highly variable genes were then scaled, and dimensionality reduction was performed using principal component analysis (PCA). Batch effects were subsequently removed using Harmony (v1.2.3)(44) implemented within Seurat.

Clustering and cell type annotations were carried out in two steps. In the first step, Louvain clustering(37) was applied using the first 30 Harmony components with a cluster resolution of 0.5. Cell classes (e.g., neurons and non-neuronal cells) were assigned to the clusters based on the relative expression of a curated list of marker genes. In the second step, cells from each cell class were isolated and sub-clustered to identify more granular clusters representing specific cell subclasses and subtypes based on a list of known marker genes (**Supplement Table 3**). This led to the identification of six non-neuronal subtypes: astrocytes, endothelial cells, microglia, oligodendrocytes, oligodendrocyte precursor cells (OPCs), and glial precursor cells (GPCs). Inhibitory neurons were the only subclass of neurons identified, and BFCNs were identified as a subtype within the inhibitory neuron population. Additional ‘low quality cells’ that could not confidently be identified with known marker genes were removed (**Supplement 1C**). After all the quality control steps and cell type annotations, the dataset retained a total of 37,467 cells and all donors were represented (**Supplement 1D-E**). There is a positive correlation (0.92) between the number of unique molecular identifiers (RNA Count) and the number of genes (RNA Features) (**Supplement 1F**).

Following separate QC and cell annotations, control and DS data were merged and batch corrected using Harmony integration in Seurat. The FindMarkers() function in Seurat was used with the MAST model and sample age as a latent variable to calculate differentially expressed genes (DEGs). For this analysis, we required that at least 25% of cells in a cluster expressed the gene for it to be considered a DEG. DEGs for each cell type are listed in **Supplement Table 4**. Hsa21 DEGs for each cell type are listed in **Supplement Table 5**. DEGs were mapped to chromosomes using the online tool, MG2C(45). Gene set enrichment analysis (GSEA) was performed using the FGSEA method(46). The scProgram and clusterProfiler(47) packages were used for KEGG pathway analysis.

### 2.6 snATAC-seq Processing and QC

The snATAC fragment data files were merged into a single master fragment file by 1) mapping the snATAC barcodes in the prepared per-sample fragment files to the corresponding snRNA seq barcodes (for direct integration), and 2) subsequently coordinate-sorting, compressing and tabulating these data for analysis in Seurat (v5.1.0)(36, 37, 48) and Signac (v1.14)(49). The multiomic integration analysis proceeded by assessing and filtering the snATAC data for those barcodes meeting the snRNA-seq quality control criteria (**Supplement G-H**).

Barcodes meeting the following criteria were selected for the integrated analysis: 1) number of fragments between 1k and 100k, 2) A TSS enrichment score >=2 and 3) nucleosome fraction score < 4 (**Supplement 1I**). These criteria resulted in 31,411 total barcodes for multiomic analysis (**Supplement 1J**).

### 2.7 Peak Calling

Peak calling was facilitated with the Signac CallPeaks() function, using MACS2 (v2.2.9.1)(50, 51). This was done for the integrated barcode data set partitioned by both annotated cell type (per the snRNA-seq analysis) and disease condition of donors (unaffected control and DS, respectively). For each cell type and condition, peaks were called, and a merged set of peak regions were compiled as implemented in CallPeaks(). A chromatin assay object representing the merged regions was aggregated by barcode and prepared in a merged Seurat object. The chief quality control measure assessed for the peak calling was the fraction of reads in peaks (FRiP) per barcode (Supplement 1K).

### 2.8 Peak-to-Gene Links

We estimated peak-to-gene links (i.e., nearby peaks correlated with gene expression of a given gene) for a few selected cell types to examine characteristics of regulatory elements across control and DS samples. These calculations were done separately across cell type and genotype. Peak-to-gene links within 500 kb of the corresponding transcription start sites (TSS) were obtained using LinkPeaks() function. The loci of genes of interest were then visualized using the coveragePlot() function in Signac, highlighting the differences in the regulatory landscape across control and DS samples.

### 2.9 Protein Carbonyl Enzyme-linked Immunosorbent Assay

15mg of frozen pulverized tissue was homogenized in 100uL of 1x PBS with 1% Protease and Phosphatase Inhibitor Cocktail (Sigma-Aldrich #PPC1010). Protein concentration was quantified with a *DC* Protein Assay following the manufacturer’s protocol (Bio-Rad #5000116). Protein oxidation was measured with a Protein Carbonyl ELISA Kit following the manufacturer’s protocol (abcam #ab238536). Samples and standards were run in duplicate and absorbance was read on a Molecular Devices VersaMax plate reader.

### 2. Malondialdehyde Lipid Peroxidation Enzyme-linked Immunosorbent Assay

15mg of frozen pulverized tissue was homogenized in 100uL of 1x PBS with 1% Protease and Phosphatase Inhibitor Cocktail (Sigma-Aldrich #PPC1010). Protein concentration was quantified with a *DC* Protein Assay following the manufacturer’s protocol (Bio-Rad #5000116). Lipid peroxidation was measured with an MDA Assay Kit (competitive ELISA) (abcam #ab238537). Protein samples were diluted to 1mg/mL and were assayed according to the manufacture’s protocol. Samples were assayed twice using fresh preparations of protein extracts for each assay run. Samples and standards were run in duplicate and absorbance was read on a Molecular Devices VersaMax plate reader. The average concentration from both assay runs were averaged.

## 3. Results

### 3.1 Overview of Human Basal Forebrain Tissue

Our data represent the first single-nucleus sequencing of the human basal forebrain. Single-nucleus multiomic analysis of gene expression and chromatin accessibility (**Figure 1B**) was performed on four unaffected control and four DS basal forebrain samples matched for age, sex, and PMI (**Figure 1A; Supplement Table 1**). After quality control (**Supplement 1A-C**), 34,467 cells were used for downstream analysis (**Figure 1C**). Cells from all donors were represented (**Supplement 1D-E**). The cell types identified in the basal forebrain were astrocytes, BFCNs, endothelial cells, inhibitory neurons, glial progenitor cells (GPCs), microglia, oligodendrocytes, and oligodendrocyte precursor cells (OPCs) (**Figure 1C**), and all cell types are present in both control and DS BF tissue samples.

Cell annotations of control and DS samples were performed separately. Initially, cells were annotated by major cell class-excitatory neurons (ExN), inhibitory neurons (InN), and non-neuronal cells (NNC). Only InNs and NNCs were identified in the initial classification (**Supplement 2A**). NNCs were sub-clustered to subclasses annotated as glial or endothelial cells (**Supplement 2B**). Astrocytes, GPCs, microglia, oligodendrocytes, and OPCs were annotated in the glial subclass (**Figure 1C; Supplement 2C**). BFCNs were identified as a subtype within inhibitory neurons (**Figure 1C; Supplement 2C**). BFCNs, which are capable of co-transmitting ACh and GABA(52, 53), were initially annotated as InNs based on their expression of *GAD1, GABBR1,* and *SLC6A1* (**Supplement 2A-B**).

After separate cluster annotation, control and DS samples were integrated and batch corrected with Harmony integration. Following integration, cell types from control and DS cluster together (**Figure 1D).** While all cell types are present in both control and DS, the proportion of each cell type differs (**Figure 1D**). Proportional to total cell number, there are fewer astrocytes and more inhibitory neurons and microglia in the DS basal forebrain (**Figure 1D**). Principal component analysis (PCA) of pseudobulk data, categorized by cell type and genotype, reveals that cell type, not genotype, has the greatest influence on gene expression between cell types (**Figure 1E**). Non-neuronal glial cells (astrocytes, GPCs, microglia, oligodendrocytes and OPCs) cluster closely together. The non-neuronal endothelial cells of the vasculature cluster separate from the other subtypes. Neurons, inhibitory and BFCNs, form a cluster (**Figure 1E**). The differences in gene expression that exist due to genotype are not enough to cause control and DS to form distinct clusters in the PCA plot. In all cell types in the basal forebrain, differentially expressed genes (DEGs) in DS compared to control are encoded across the genome (**Figure 1F; Supplement 2D**), consistent with other gene expression data from DS samples(54–61).

Next, we assessed differences in chromatin accessibility and peak-to-gene linkages of Hsa21-encoded genes that could account for differential gene expression in the DS basal forebrain. Following QC of the ATAC-seq data (**Supplement 1G-I**), 16,771 control cells and 14,640 DS cells were used for downstream analysis (**Supplement 1J**). The integrated multiomic data was clustered, revealing that all cell types passed ATAC QC and were present in both control and DS samples (**Figure 1G**).

### 3.2 Hsa21-Encoded Genes

We investigated whether and how Hsa21-encoded genes are dysregulated in DS. Of the 221 predicted Hsa21 protein-coding genes annotated in the GRCh38.p14 reference assembly, 85 Hsa21 genes are dysregulated between astrocytes, BFCNs, inhibitory neurons, microglia, and oligodendrocytes, with most genes dysregulated in more than one cell type (**Figure 2A; Supplement Table 5**). Hsa21 genes represent 1.7% of astrocyte DEGs, 10.6% of BFCN DEGs, 2.0% of inhibitory neuron DEGs, 3.1% of microglia DEGs, and 2.0% of oligodendrocyte DEGs (**Figure 2B; Supplement 3A-E**). These dysregulated Hsa21 genes account for 26.7%, 4.1%, 33.0%, 10.9%, and 21.3% of total Hsa21 protein-coding genes, respectively (**Figure 2C**). 56 Hsa21 genes are upregulated and 3 are downregulated in DS astrocytes (**Figure 2A, 2D**); 9 are upregulated in DS BFCNs (**Figure 2A, 2E**); 72 are upregulated and 1 is downregulated in DS inhibitory neurons (**Figure 2A, 2F**); 21 are upregulated and 3 are downregulated in DS microglia (**Figure 2A, 2G**); 42 are upregulated and 5 are downregulated in DS oligodendrocytes (**Figure 2A, 2H**). 79 Hsa21-encoded genes are dysregulated between endothelial cells, GPCs, and OPCs (**Supplement 3F–I**), accounting for 2.4%, 3.1%, and 1.8% of total DEGs in these cells (**Supplement 3J**). Hsa21-encoded genes account for 22.2%, 24.9%, and 22.6% of total Hsa21 protein-coding genes, respectively (**Supplement 3K-N**). Gene set enrichment analysis (GSEA) of biological processes reveals that upregulated Hsa21 genes in all cell types are enriched in regulation of developmental process, neuron differentiation, and neurogenesis (**Figure 2I**). These results reveal dysregulated Hsa21 genes in the basal forebrain that may contribute to the heterochronic development reported in DS(62–64).

**Figure 2.**
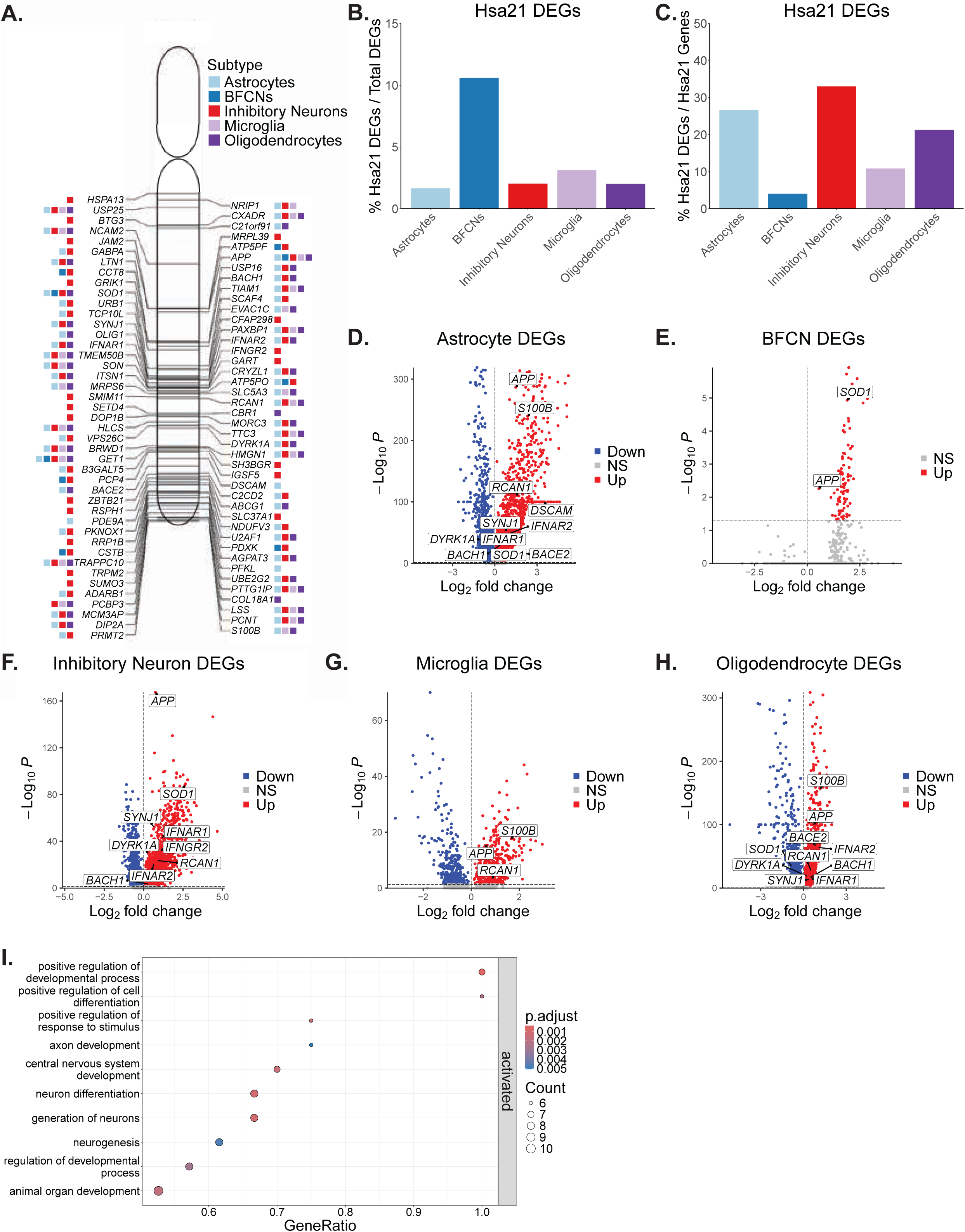
A) Hsa21-encoded genes differentially expressed in astrocytes, BFCNs, inhibitory neurons, microglia, and oligodendrocytes. Boxes next to each gene represent the cell types that gene is dysregulated in. B) Percent of Hsa21 DEGs relative to total DEGs per cell type. C) Percent of Hsa21 DEGs normalized to total Hsa21 protein-coding genes per cell type. D) Volcano plot of dysregulated genes in DS astrocytes with Hsa21 genes of interest labeled. E) Volcano plot of dysregulated genes in DS BFCNs with Hsa21 genes of interest labeled. F) Volcano plot of dysregulated genes in DS inhibitory neurons with Hsa21 genes of interest labeled. G) Volcano plot of dysregulated genes in DS microglia with Hsa21 genes of interest labeled. H) Volcano plot of dysregulated genes in DS oligodendrocytes with Hsa21 genes of interest labeled. Hsa21 genes of interest that are labeled on the volcano plots are *APP, S100B, DYRK1A, DSCAM, SOD1, RCAN1, IFNAR1, IFNAR2, IFNGR2, SYNJ1, BACE2,* and *BACH1*. I) Gene set enrichment analysis of Hsa21 genes dysregulated across all subtypes. Dysregulated Hsa21 genes are enriched in regulation of developmental process, neuron differentiation, and neurogenesis processes.

To further investigate the impact of the additional copy of Hsa21 in basal forebrain cells, we analyzed the chromatin accessibility and peak-to-gene linkages of differentially expressed Hsa21 genes in astrocytes, BFCNs, inhibitory neurons, microglia, and oligodendrocytes. However, the number of BFCNs that passed ATAC QC was too low to calculate statistically significant peak-to-gene linkages in these cells. Two genes per cell subtype were selected from the Hsa21 DEGs to calculate peak-to-gene linkages for the other subtypes (**Supplement 4**). In DS astrocytes and microglia, larger peaks are present at the transcription start site (TSS) of *APP* and *S100B* (**Supplement 4A, 4C**). DS inhibitory neurons have larger peaks at the TSS of *APP* and *SOD1* (**Supplement 4B**). TSS peaks in *APP* and *OLIG1* are increased in DS oligodendrocytes (**Supplement 4E**). Since peaks mark regions of accessible chromatin, these results suggest increased chromatin accessibility, facilitating the recruitment of transcription factors and enhancers, which in turn promote active gene transcription. The peak-to-gene links observed in *APP* and *S100B* (DS astrocytes), *APP* (DS inhibitory neurons), *APP* and *S100B* (DS microglia), and *OLIG1* (DS oligodendrocytes) may provide insight into potential gene regulatory elements that control expression of these genes in DS (**Supplement 4A-D**). Future work identifying these potential regulatory motifs and determining their necessity for gene regulation will provide insight into the mechanisms controlling the expression of these genes in DS.

### 3.3 Astrocytes

Recently, the important role of glia in neurodegenerative diseases, including DS(65, 66), has emerged. The proportion of astrocytes is decreased and the proportion of microglia is increased in DS relative to total glial cells (**Supplement 5A**) and total cells (**Figure 1D**). The differential proportions of these cells may have functional consequences on both development and degeneration in the BF.

We analyzed the differential gene expression in astrocytes, microglia, and oligodendrocytes to identify signatures of early deficits in glial cells of the DS basal forebrain. Identification of DEGs in DS astrocytes (adjusted *P*<.05) revealed dysregulated genes across the genome(**Figure 3A; Supplement Table 4**). Hsa21 genes constitute only 1.7% of the dysregulated genes in astrocytes (**Figure 2B**). More Hsa21 genes are represented than all other chromosomes except Hsa13 and Hsa18 in DS astrocytes when normalizing the DEGs per chromosome to the number of protein-coding genes of the chromosome (**Figure 3B**). The top 30 DEGs (from the largest absolute values of the LFCs) in DS astrocytes are all upregulated genes (**Figure 3C**) GSEA of biological processes reveals that upregulated genes in DS astrocytes are enriched in ribosome biogenesis, translation, and ensheathment of neurons and axons (**Figure 3D**). Meanwhile, downregulated genes are enriched in protein folding and protein maturation (**Figure 3D**). The five most dysregulated pathways in DS astrocytes-ribosome biogenesis, cytoplasmic translation, translation, ensheathment of neurons, and axon ensheathment-consist of genes that are upregulated are shown in the gene-concept network plot (cnet plot) (**Figure 3E; Supplement Table 6**). The dysregulation of ribosome and translation genes and genes involved in neuron ensheathment suggests that DS astrocytes are increasing the production of proteins that mediate interactions with neurons in the BF.

**Figure 3.**
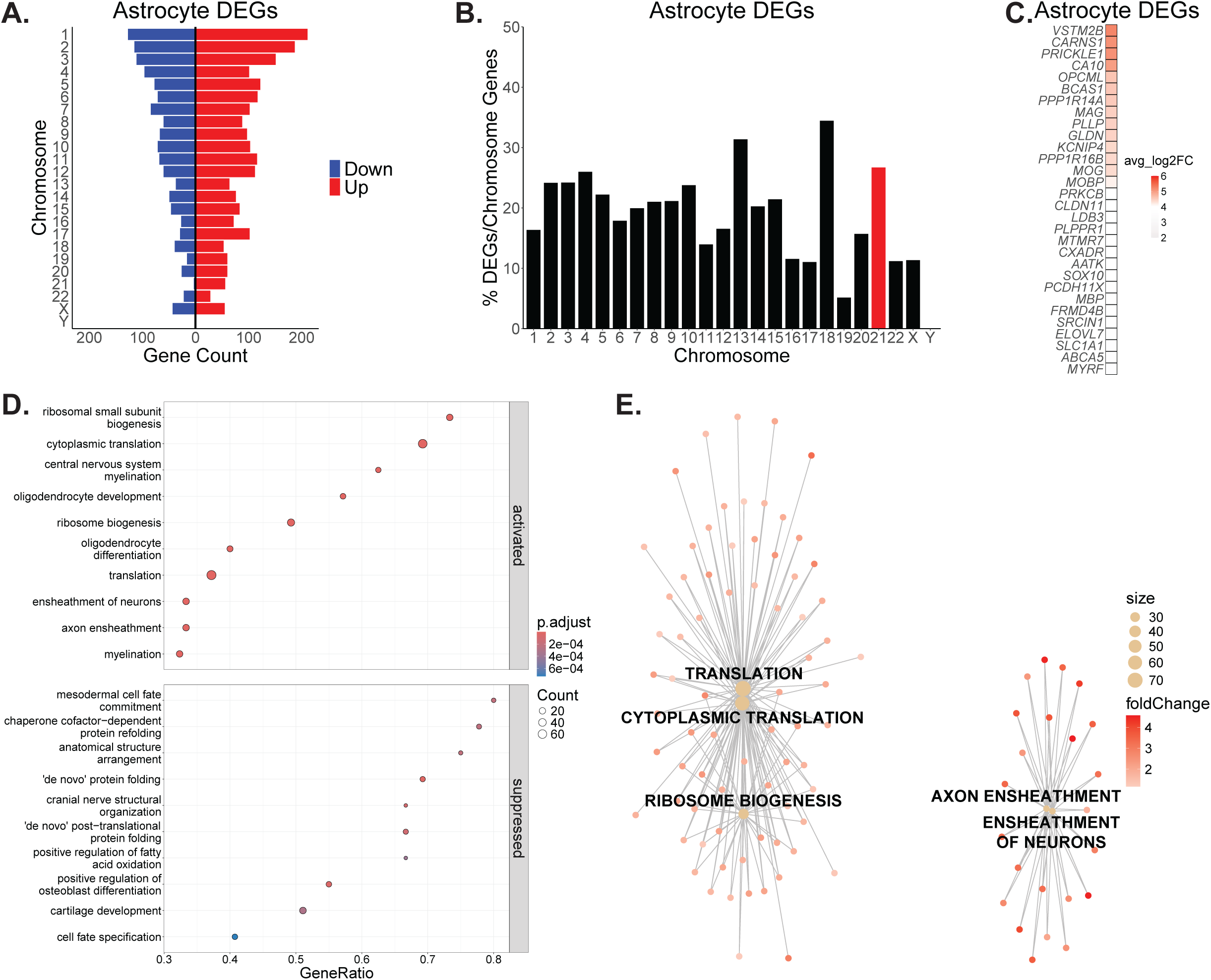
A) DEGs are distributed across the genome in DS astrocytes. B) Percent of DEGs normalized to protein-coding genes per chromosome in DS astrocytes with Hsa21 highlighted in red. C) Top 30 DEGs in DS astrocytes. D) DS astrocyte upregulated genes are enriched in ribosome biogenesis, translation, and ensheathment of neurons and axons while downregulated genes are enriched in protein folding and protein maturation. E) CNET plot of the top 5 dysregulated biological processes in DS astrocytes. Genes for each category node are listed in Supplement Table 6.

### 3.4 Microglia

DEGs (adjusted *P*<.05) in DS microglia are encoded across the genome(**Figure 4A; Supplement Table 4**). Hsa21-encoded genes are represented at a higher percentage in DS microglia when taking into account the number of protein-coding genes per chromosome (**Figure 4B**). The top 30 DEGs in DS microglia comprised of 20 upregulated genes and 10 downregulated genes (**Figure 4C**). GSEA reveals that upregulated genes in DS microglia of the basal forebrain are enriched in complement activation and translation whereas downregulated genes are enriched in transcription, protein folding, and protein maturation (**Figure 4D**). The five most dysregulated pathways in DS basal forebrain microglia are upregulated complement activation and cytoplasmic translation processes. Nucleic acid metabolic process, protein maturation processes are downregulated in DS basal forebrain microglia (**Figure 4E; Supplement Table 6**). These results reveal that, at birth, there is evidence of an immune response via activation of the complement system and dysregulation of microglial function in the DS BF.

**Figure 4.**
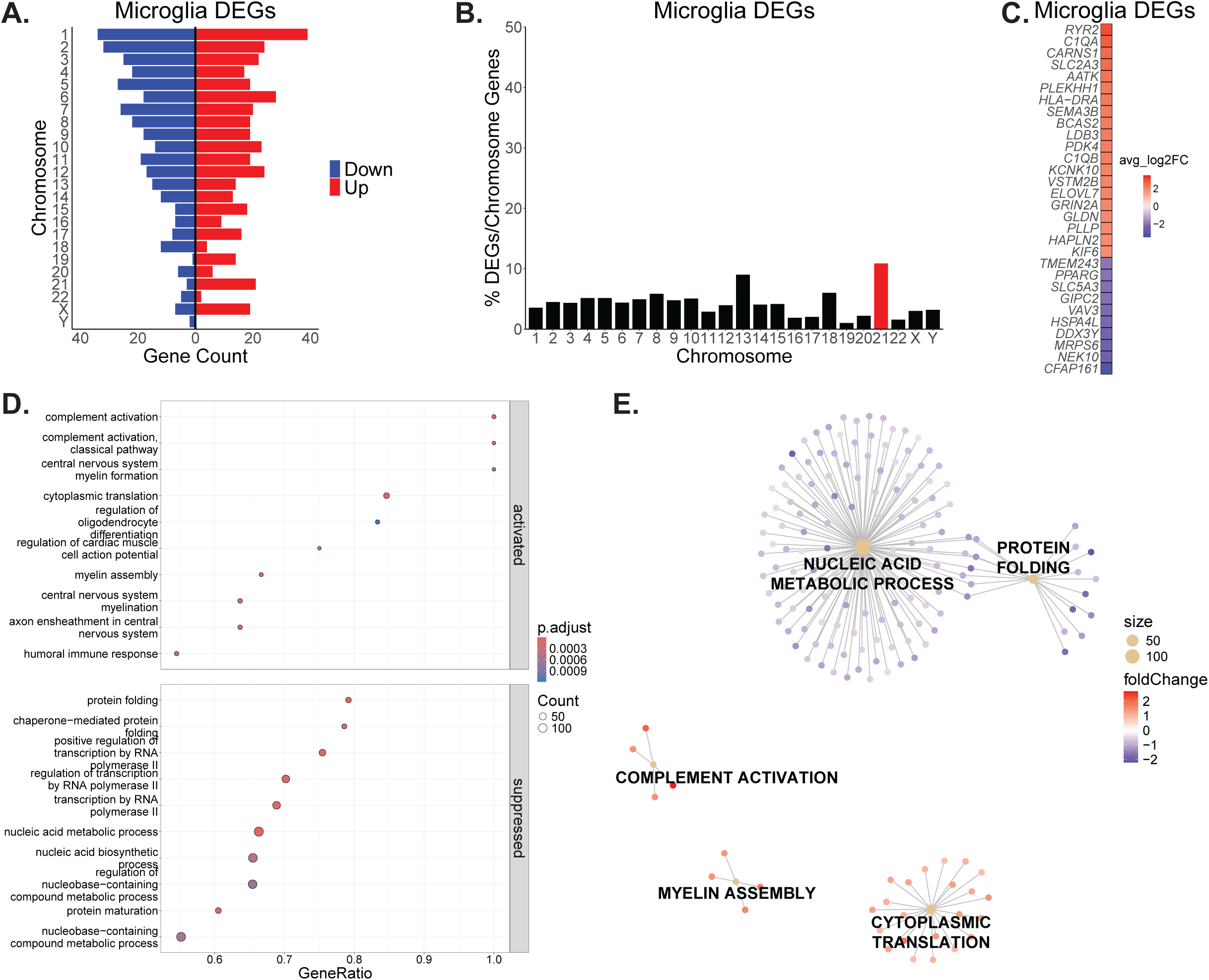
A) DEGs are distributed across the genome in DS microglia. B) Percent of DEGs normalized to protein-coding genes per chromosome in DS microglia with Hsa21 highlighted in red. C) Top 30 DEGs in DS microglia. D) DS microglia upregulated genes are enriched in complement activation and translation whereas downregulated genes are enriched in transcription, protein folding, and protein maturation. E) CNET plot of the top 5 dysregulated biological processes in DS microglia. Genes for each category node are listed in Supplement Table 6.

### 3.5 Oligodendrocytes

In oligodendrocytes, DEGs (adjusted *P*<.05) are encoded across the genome(**Figure 5A; Supplement Table 4**). Similar to the other glial cells, Hsa21 genes are represented at a higher percentage in DS oligodendrocytes when normalizing DEGs to the chromosome’s number of protein-coding genes (**Figure 5B**). The top 30 DEGs include 28 downregulated and 2 upregulated genes in DS oligodendrocytes (**Figure 5C**). Upregulated genes in DS oligodendrocytes are enriched in metabolic and secretion processes (**Figure 5D)**. Downregulated genes are enriched in protein folding and response to stimuli processes (**Figure 5D**). The top five dysregulated processes-response to temperature stimulus, response to heat, chaperone-mediated protein folding, ‘de novo’ protein folding, and protein folding- are all downregulated in in DS oligodendrocytes of the basal forebrain (**Figure 5E; Supplement Table 6**). These results suggest that DS oligodendrocytes may have an increase in protein misfolding and may not be as responsive to environmental cues, potentially contributing to the reduced myelination reported in DS (56, 67, 68).

**Figure 5.**
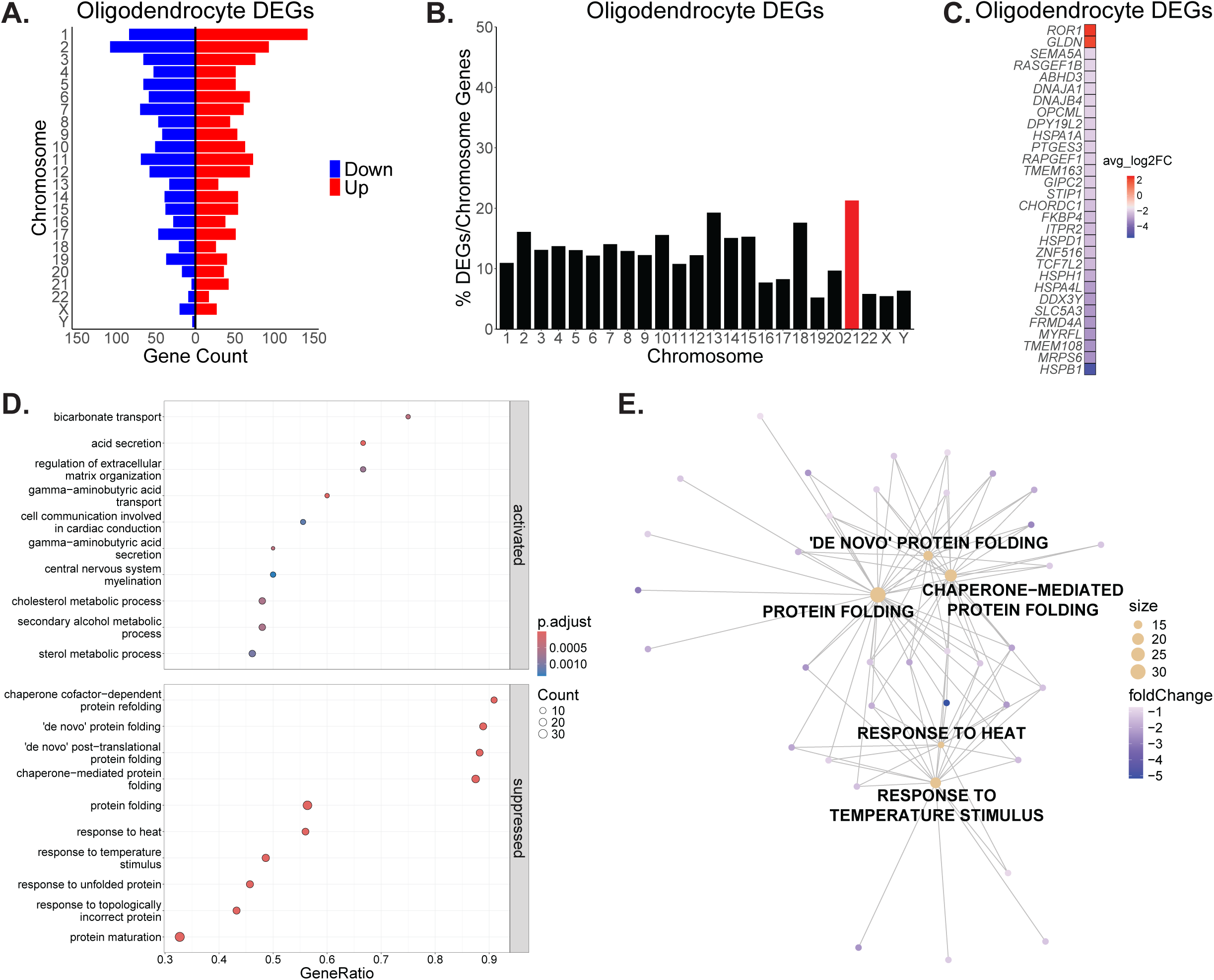
DEGs are distributed across the genome in DS oligodendrocytes. B) Percent of DEGs normalized to protein-coding genes per chromosome in DS oligodendrocytes with Hsa21 highlighted in red. C) Top 30 DEGs in DS oligodendrocytes. D) DS oligodendrocyte upregulated genes are enriched in metabolic and secretion processes, and downregulated genes are enriched in protein folding and response to stimuli processes. E) CNET plot of the top 5 dysregulated biological processes in DS oligodendrocytes. Genes for each category node are listed in Supplement Table 6.

### 3.6 Inhibitory Neurons

Inhibitory neurons account for 9.5% of control cells and 34.1% of DS cells in our data (**Figure 1D; Supplement 2A-B**). DS inhibitory neurons have DEGs (adjusted *P*<.05) distributed across the genome (**Figure 6A; Supplement Table 4**). Hsa21 genes make up a small percentage of DS inhibitory DEGs (**Figure 2B**) but are overrepresented when normalizing the DEGs per chromosome to the number of chromosome protein-coding genes (**Figure 6B**). The top 30 DEGs in DS inhibitory neurons based on the absolute value of the LFC are all upregulated (**Figure 6C**). GSEA reveals that upregulated genes in DS Inhibitory neurons of the basal forebrain are enriched in processes of translation, ATP synthesis, and energy metabolism (**Figure 6D**). Downregulated genes are enriched in cell adhesion and synaptic transmission pathways (**Figure 6D**). The top five dysregulated pathways- ribosomal small subunit biogenesis, cytoplasmic translation, proton transmembrane transport, ATP synthesis coupled electron transport, and oxidative phosphorylation- are upregulated processes in DS Inhibitory neurons of the basal forebrain (**Figure 6E; Supplement Table 6**). DS Inhibitory neurons display dysregulation of genes associated with ATP production, cell adhesion, and synapse formation, suggesting mitochondrial defects and cell-cell communication deficits may already be present at birth in the DS basal forebrain.

**Figure 6.**
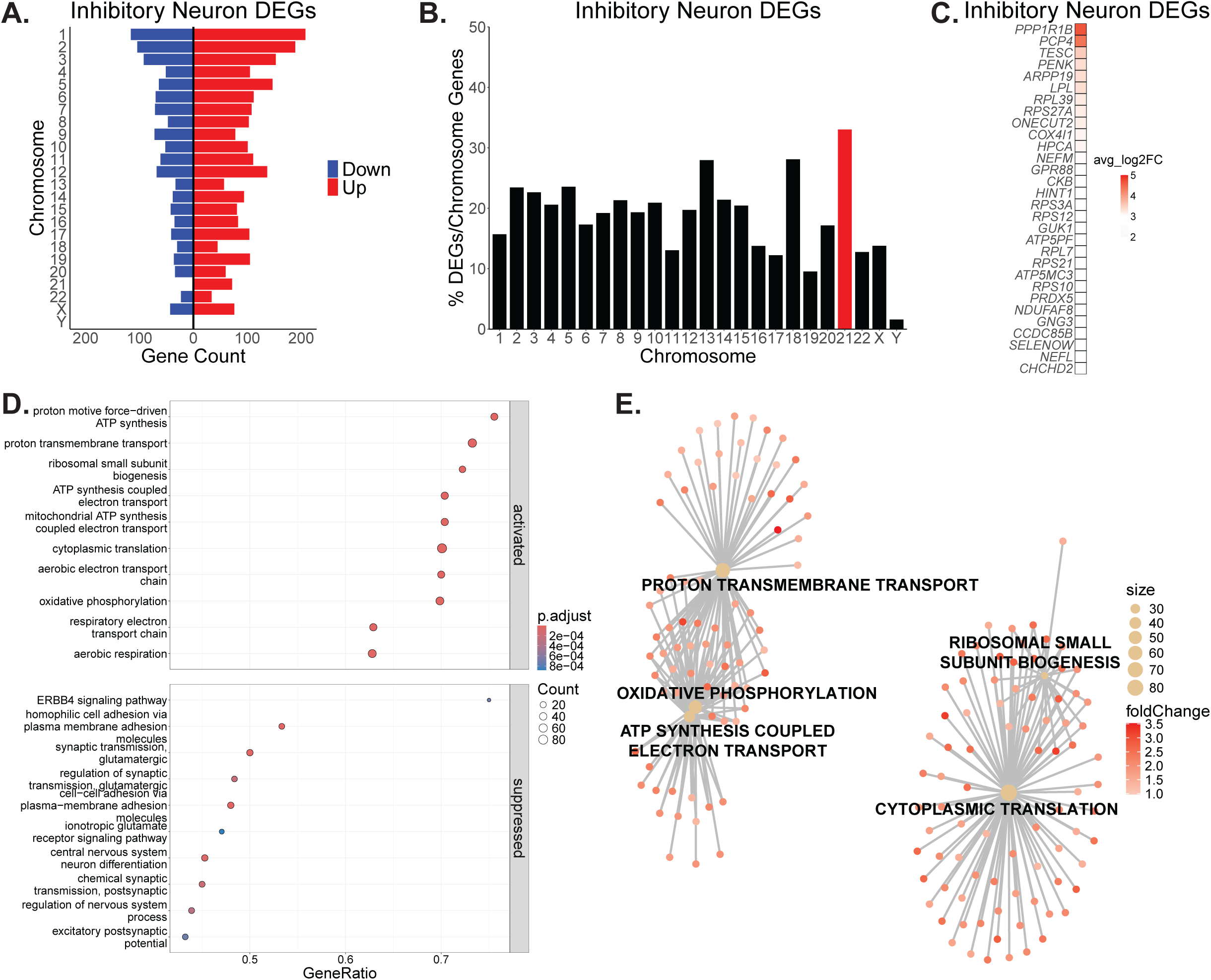
A) DEGs are distributed across the genome in DS inhibitory neurons. B) Percent of DEGs normalized to protein-coding genes per chromosome in DS inhibitory neurons with Hsa21 highlighted in red. C) Top 30 DEGs in DS inhibitory neurons. D) DS inhibitory neuron upregulated genes are enriched in processes of translation, ATP synthesis, and energy metabolism. Downregulated genes are enriched in cell adhesion and synaptic transmission pathways. E) CNET plot of the top 5 dysregulated biological processes in DS inhibitory neurons. Genes for each category node are listed in Supplement Table 6.

### 3.7 Two Distinct BFCN Populations

To identify the BFCNs for downstream analysis, we sub-clustered the inhibitory neurons and annotated BFCN clusters based on known markers (**Figure 7A; Supplement Table 3**). Of the total inhibitory neurons in control and DS, BFCNs comprise 5.2% and 2.4%, respectively (**Figure 7A**). snRNA-seq provides an unbiased approach to identify potential novel markers for BFCNs. However, relative to inhibitory neurons, many of the top BFCN marker genes have already been characterized in cholinergic neurons (**Supplement 5B**).

**Figure 7.**
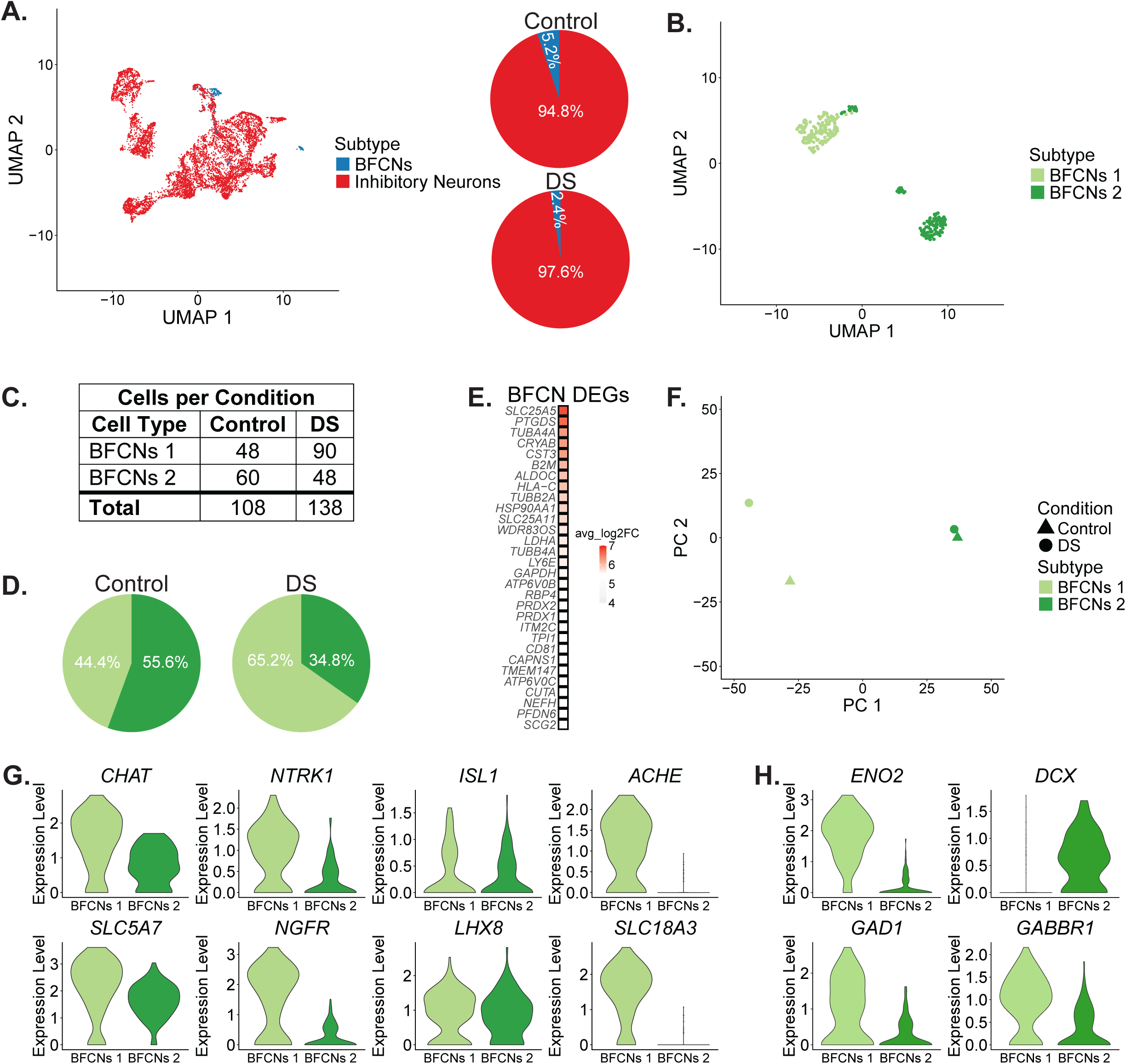
A) UMAP of all inhibitory neurons. BFCNs cluster in two distinct clusters. Proportion of BFCN subpopulations in control and DS. B) BFCNs were subset from the data and reclustered, again clustering into two distinct populations that were labeled BFCNs 1 and BFCNs 2. C) Counts of BFCNs1 and BFCNs per condition. D) Proportion of BFCN populations in control and DS. E) BFCNs 1 DEGs relative to BFCNs 2. F) PCA of BFCN populations. There is more variability between control and DS BFCNs 1. G) Expression of known cholinergic marker genes in BFCN subpopulations. BFCNs 2 have lower expression of most of these genes. H) Expression of immature and mature neuron marker genes in BFCN subpopulations. BFCNs 1 have higher expression of *ENO2*, a mature neuron marker, while BFCNs 2 have increased expression of DCX, an immature neuron marker. BFCNs 1 also have higher expression of *GAD1*, encoding the enzyme that catalyzes the conversion of glutamate into GABA, and *GABBR1*, a GABA receptor

Interestingly, BFCNs formed 2 separate cluster (**Figure 7A**). We then subset the BFCNs and sub-clustered them, again resulting in the separation of two distinct clusters (**Figure 7B**). The two clusters were annotated as BFCNs 1 and BFCNs 2 (**Figure 7B**). In controls these two populations of BFCNs are roughly of equal proportion, whereas in DS there are more BFCNs 1 (**Figure 7C-D**). Differentially expressed genes calculated between BFCNs 1 and BFCNs 2 reveal that several tubulin genes and several genes involved in energy metabolism are upregulated in BFCNs 1 (**Figure 7E**). PCA of the BFCN subpopulations shows that there is greater variability in the BFCNs 1 population (**Figure 7F).** These results suggest that BFCNs 1 are more impacted by DS than BFCNs 2.

We compared the expression of known BFCN markers to identify differences in these two BFCN populations. Both BFCN populations express the established marker genes of BFCNs, including genes encoding enzymes and transporters in the acetylcholine (ACh) pathway (*CHAT, ACHE*, *SLC18A3, SLC5A7)*, neurotrophic receptors required for the maintenance and survival of BFCNs (*NTRK1, NGFR*), and transcription factors that regulate these genes (*ISL1, LHX8*). However, BFCNs 2 express most of these genes at lower levels compared to BFCNs 1 (**Figure 7G**). BFCNs 2 have a significant reduction in the expression of *SLC18A3,* a transmembrane protein responsible for transporting ACh into secretory vesicles for release, and *ACHE*, which hydrolyzes ACh into choline that is recycled for continued ACh synthesis (**Figure 7G**). These results suggest that BFCNs 2, with reduced expression of essential components for ACh neurotransmission, are likely not fully functional cholinergic neurons.

Given the developmental stage of the samples and the reduced expression of genes in the ACh pathway, we suspected that BFCNs 2 are immature compared to BFCNs 1. We assessed the expression of mature and immature neuron markers in these two populations (**Figure 7H**). Compared to BFCNs 1, BFCNs 2 have decreased expression of *ENO2*, a mature neuron marker, and increased expression of *DCX*, an immature neuron marker (**Figure 7H**). Additionally, BFCNs 2 exhibit decreased expression of the GABA receptor, *GABBR1,* along with decreased *GAD1* expression, which encodes GAD65, the enzyme that catalyzes the conversion of glutamate into GABA (**Figure 7H**). GSEA reveals that energy metabolism pathways are activated in BFCNs 1 whereas cell adhesion and processes related to synaptic formation are suppressed (**Supplement 5C; Supplement Table 6**). KEGG pathway analysis reveals upregulated genes in BFCNs 1 are enriched in oxidative phosphorylation and glycolysis/gluconeogenesis pathways (**Supplement 5D**). GSEA and KEGG analysis suggests that BFCNs 1 are likely more metabolically active, characteristic of more mature neurons. This increased metabolic activity coupled with the increased expression of genes in the ACh and GABA pathways, the increased expression of *ENO2*, and the decreased expression of *DCX* suggest BFCNs 1 are a more mature population of cells relative to BFCNs 2. The increased proportion of mature BFCNs 1 in DS supports precocious development of DS BFCNs (**Figure 7D**). The limited number of DEGs in DS BFCNs 2 (**Supplement 5E; Supplement Table 4**) precludes pathway analysis and so further analysis was performed only on the BFCNs 1 population.

### 3.8 Analysis of Mature BFCNs

We analyzed the DEGs (adjusted *P*<.05) of BFCNs 1 to understand the cellular mechanisms that may contribute to DS BFCN degeneration so early in life. Differentially expressed genes in BFCNs are distributed across the genome (**Figure 8A**). DS BFCNs 1 have 89 upregulated genes and six downregulated genes relative to control (**Figure 8A; Supplement Table 4**). Hsa21 genes make up 10.6% of DS BFCNs 1 DEGs (**Figure 2B**) but are overrepresented when normalizing the DEGs per chromosome to the number of chromosome protein-coding genes (**Figure 8B**). Of the top 30 DEGs in DS BFCNs 1, all are upregulated (**Figure 8C; Supplement Table 4**). Genes upregulated in DS include those encoding subunits of the oxidative phosphorylation pathway (*NDUFS2, COX5A, ATP5PO, ATP5PF,* and *UQCRC2),* antioxidant enzymes (*SOD1, PRDX1,* and *PRDX2)*, and subunits of the vacuolar-type ATPase (V-ATPase) (*ATP6V0B, ATP6V0D1,* and *ATP6V0C)* (**Figure 8C; Supplement Table 4**). Overexpression of V-ATPase, an ATP-driven proton pump that regulates cellular pH and plays a role in overall cell homeostasis, has been linked to several human diseases(69). Additionally, genes associated with glycolysis (*PGAM1, LDHB, LDHA, PDHA1, TPI1, GAPDH,* and *PGK1*) are also upregulated in DS BFCNs (**Figure 8C; Supplement Table 4**). The increased expression of glycolysis-associated genes in neurons that should be relying primarily on the oxidative phosphorylation pathway for ATP production suggests DS BFCNs 1 may be starting to shift toward glycolysis as the primary source for energy production, a shift that is linked to mitochondrial dysfunction and several neurodegenerative diseases(70). Although several genes related to energy metabolism are dysregulated in DS BFCNs, potential compensatory mechanisms may also be at play, evidenced by the increased expression and increased chromatin accessibility of antioxidant enzymes (**Supplement 5F**) as well as the increased expression of V-ATPase subunits, all of which help maintain cellular homeostasis.

**Figure 8.**
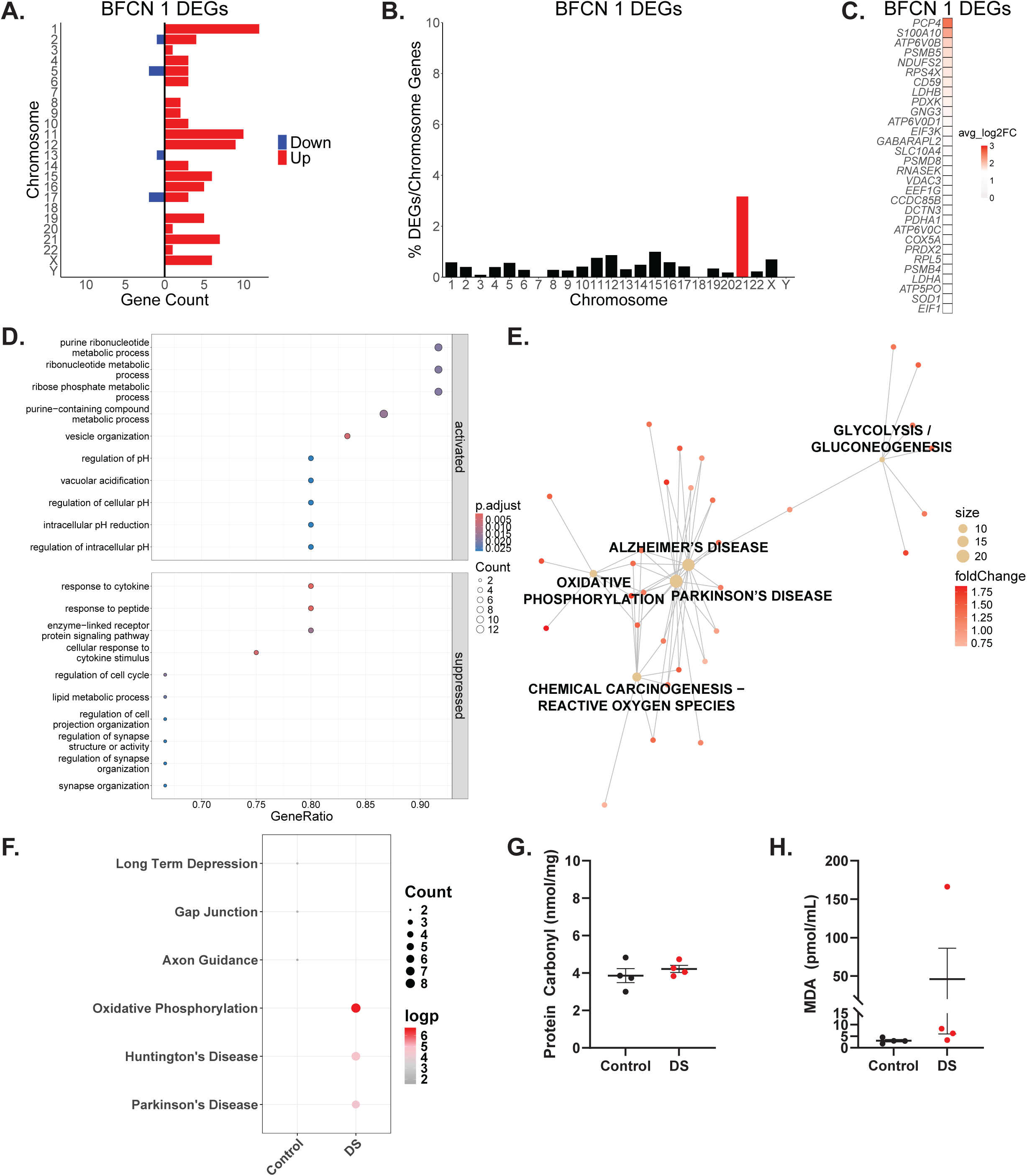
A) DEGs are distributed across the genome in DS BFCNs 1. B) Percent of DEGs normalized to protein-coding genes per chromosome in DS BFCNs 1 with Hsa21 highlighted in red. C) Top 30 DEGs in DS BFCNs 1. D) Metabolic and regulation of pH processes are activated in DS BFCNs 1. Response to stimuli and synapse formation processes are suppressed in DS BFCNs 1. E) CNET plot of 5 dysregulated KEGG pathways in DS BFCNs 1 calculated with clusterProfiler. Dysregulated BFCNs 1 genes are enriched oxidative phosphorylation and glycolysis/gluconeogenesis pathways and are associated with several neurodegenerative diseases. Genes for each category node are listed in Supplement Table 6. F) KEGG pathway analysis of enriched genes in control and DS BFCNs 1 calculated with scProgram. In DS BFCNs 1, DEGs are enriched in the oxidative phosphorylation pathway and these DEGs are associated with Parkinson’s and Huntington’s diseases. G) Protein oxidation assay from the bulk basal forebrain tissue. There is no difference in the protein carbonyl content in control and DS, indicating no change in protein oxidation in the DS basal forebrain. H) Lipid peroxidation assay from the bulk basal forebrain tissue. There is an increase in the malondialdehyde (MDA) content in DS, indicating increased lipid peroxidation in the DS basal forebrain. The large variability between DS samples suggests that ROS are just beginning to accumulate in the DS basal forebrain during this period of development.

GSEA of biological processes reveals that metabolic and regulation of pH processes are activated in DS BFCNs 1 of the basal forebrain (**Figure 8D**). Response to stimuli and synapse formation processes are suppressed in DS BFCNs 1 (**Figure 8D**). Gene-concept network analysis of KEGG pathway enrichment performed with clusterProfiler reveals that, in addition to oxidative phosphorylation and glycolysis/gluconeogenesis pathways, genes upregulated in DS BFCNs 1 are enriched in pathways associated with several neurodegenerative diseases, including AD, PD, Huntington’s disease, prion disease, and amyotrophic lateral sclerosis (ALS) (**Figure 8E; Supplement Table 6**). KEGG enrichment analysis performed with scProgram reveals that DS BFCNs 1 DEGs are enriched in the oxidative phosphorylation pathway (**Figure 8F**), further suggesting that there is metabolic dysregulation in DS BFCNs 1.

Gene expression and chromatin accessibility of antioxidant genes indicates there may be an increase in OXPHOS and resulting ROS in DS BFCNs 1. To validate this finding, we probed for lipid peroxidation and protein oxidation, downstream consequences of ROS, to determine if ROS is accumulating in the DS basal forebrain at birth. Protein carbonylation, the oxidation of protein side chains induced by ROS, is not altered in the DS basal forebrain (**Figure 8G**). Though variable and not statistically significant, there is an increase in lipid peroxidation in the DS basal forebrain (**Figure 8H**). Changes in lipid peroxidation but not in protein oxidation in DS samples suggests that ROS are just beginning to accumulate and induce damage to lipids, which are more susceptible to ROS(71–73), in the DS basal forebrain. Altogether, these results suggest that metabolic dysregulation and ROS are present in the DS basal forebrain at birth, potentially contributing to the early degeneration of DS BFCNs(14).

## 4. Discussion

### 4.1 Summary of findings

Our results provide the first gene expression and chromatin accessibility data of the human basal forebrain from either healthy or diseased individuals. Cholinergic neurons in the basal forebrain are vulnerable in age-associated degeneration and implicated in the progression of multiple neurodegenerative diseases, including AD, PD, and DLB(74). BFCN degeneration is particularly early in the progression of DS-AD, detectable when individuals are in their twenties(14), so these data establish a rich resource for further investigation of neurodegeneration. Thus, our findings of early neuropathology in both neurons and glia of the basal forebrain in DS may provide insight into shared mechanisms between several neurodegenerative disorders.

### 4.2 Limitations of tissue analysis

All studies analyzing post-mortem human tissue suffer from specific features of the donor and tissue including cause of death, post-mortem interval, and sample integrity that can affect the results. Samples used in this study were from donors with varied causes of death (**Supplement Table 1**) and so transcriptional changes due to specific causes of death are unlikely. Analyzed samples are from early postnatal donors, but we lack information of gestational duration. We cannot rigorously match samples for stages of development and analyzed samples that correspond to neonatal and very early infancy, a developmental phase that includes rapid growth, synaptic remodeling and myelination(75).

The organization of cholinergic neurons into four nuclei (Ch1-4) whose anatomical boundaries are not discrete(76, 77) limits our ability to determine in which nucleus or nuclei the cholinergic neurons we analyzed reside. However, degeneration of the anteromedial basal forebrain (Ch1-3) and posterior basal forebrain (Ch4) occurs concomitantly in DS(14), so any changes likely apply to BFCNs from all nuclei.

### 4.3 Single cell transcriptomic signature of Early Postnatal Basal Forebrain

All expected cell types are present in both control and DS early postnatal DS basal forebrain, but cell type proportions are altered in DS. Interestingly, the proportions of neurons and glia are contrary to reports in the DS cortex, where neurons are reduced, and astrocytes are increased compared to controls(62, 64, 78). These results suggest that prenatal development of the basal forebrain may be altered in DS, leading to the generation of different numbers of progenitors and/or neurons or, alternatively, that neuron degeneration begins prenatally. Analysis of prenatal DS tissue and induced pluripotent stem cell models are needed to interrogate earlier developmental time periods to test these possibilities.

We identified molecular events that occur in DS prior to cholinergic dysfunction(28–30) that provide clues to the vulnerability of BFCNs. We uncovered dysregulation of genes in all cell types in the early postnatal DS basal forebrain. Few dysregulated genes and molecular pathways were shared across cell types, suggesting that the gene expression differences in DS are largely cell type-specific. Although Hsa21 genes were overrepresented when normalizing to the protein-coding genes, Hsa21-encoded genes were a small proportion of total dysregulated genes in all cell types. Functional validation of these gene expression differences will define how each cell type is affected in DS.

We uncovered two populations of BFCNs (BFCNs 1, BFCNs 2) in the early postnatal forebrain that are present in control and DS samples. Both populations express established cholinergic marker genes, although BFCNs 2 express these genes at much lower levels. Additionally, BFCNs 2 have increased expression of the immature neuron marker *DCX*, suggesting that these cells are less mature and not fully functional. It is likely, given the early age of the samples, that BFCNs are still developing, resulting in immature and mature populations. However, the DS samples have a higher proportion of the BFCNs 1 population. If the BFCNs 1 population indeed represent mature neurons, then these data align with the idea of developmental heterochrony that has been proposed in DS(62–64), in which development progresses precociously in DS.

### 4.4 BFCN Degeneration

Despite the fact that BFCNs degenerate earlier in DS than in other neurodegenerative disorders (14), there is no analysis of the human DS basal forebrain in early life. Previous studies have examined the basal forebrain cholinergic system in DS and DS-AD, but the youngest individuals in these studies are adolescents and young adults(11, 14) where AD pathology has already begun (26, 27). We sought to identify molecular signatures defining vulnerability that occur in DS prior to cholinergic dysfunction(28–30). Our results reveal that basal forebrain pathology is present as early as birth in DS, indicating pathological processes begin prenatally. These results suggest that the perinatal period is a potential window of therapeutic opportunity to mitigate emerging neuropathology in the DS basal forebrain.

Mitochondrial dysfunction and dysregulated energy metabolism are emerging as hallmarks of many neurodegenerative diseases, including those that include BFCN degeneration(70, 79). Relative to control, the more mature population of DS BFCNs (BFCNs 1) upregulate several genes that encode components of the OXPHOS pathway, along with antioxidant enzymes, *PRDX1, PRDX2,* and *SOD1*, that detoxify reactive oxygen species (ROS) byproducts generated from OXPHOS. In response to ROS accumulation, NRF2 is activated which regulates the expression of antioxidant enzymes, including superoxide dismutases and peroxiredoxins(80). The upregulation of these antioxidant enzymes is potentially a compensatory mechanism to detoxify excessive ROS levels in the DS basal forebrain. We hypothesize that this early dysregulation of the OXPHOS pathway leads to an accumulation of ROS and the resulting oxidative stress increases the vulnerability of DS BFCNs. Our gene expression and biochemical assay results suggest that dysregulated energy metabolism and the early accumulation of ROS in the DS basal forebrain underlies the susceptibility of BFCNs to degeneration later in life.

While our results indicate that dysregulated genes in DS BFCNs are associated with several neurodegenerative diseases, OXPHOS genes are typically downregulated with age and in neurodegenerative diseases as cells shift toward glycolysis as the primary source for ATP production, a shift known as the Warburg effect(70, 81). Several nuclear-encoded OXPHOS subunits are upregulated in DS BFCNs, suggesting that the OXPHOS pathway is still utilized at birth in DS. In the Ts65Dn mouse model of DS and AD, OXPHOS genes are downregulated in BFCNs at 6 months of age(82, 83), approximately when BFCN dysfunction and degeneration begins in this model(84, 85). The upregulation of OXPHOS genes in early postnatal human BFCNs and the downregulation in Ts65Dn 6-month BFCNs supports this potential shift in energy metabolism as these neurons begin to degenerate. The increase in genes encoding glycolytic enzymes suggests DS BFCNs may be in the early stages of shifting toward glycolysis as the primary source for energy production. A prolonged shift from OXPHOS to glycolysis can create an energy deficit that makes cells more susceptible to oxidative stress and cell death(70). The accumulation of ROS coupled with a shift from OXPHOS to glycolysis by birth may be an early driver of BFCN vulnerability in DS.

Alternatively, the upregulation of genes encoding antioxidant enzymes and components of the glycolysis pathway may be attributed to the predominance of female samples in our study. Biological sex influences the progression of DS-AD pathogenesis(86), and sex differences have been reported in the basal forebrain cholinergic system of the Ts65Dn mouse model of DS and AD(87). Recent spatial transcriptomic analyses of DS-AD samples reveal that genes involved in oxidative stress and glucose metabolism are upregulated in females compared to males(88). The upregulation of genes encoding antioxidant enzymes, the primary defense against oxidative stress, and components of the glycolysis pathway in our study may result from the fact that three out of four control and DS samples are from female donors. However, our study is not sufficiently powered to assess the impact of sex differences on the transcriptome of DS BFCNs.

### 4.5 Non-neuronal cell dysregulation

Differential gene expression in astrocytes, microglia, and oligodendrocytes indicates neuroinflammation and myelination deficits are present in the DS basal forebrain as early as birth. DS astrocytes and microglia upregulate genes involved in translation and are potentially altering their proteomic profiles as they transition from a homeostatic state to a reactive or activated state, respectively. Activation of the complement system further suggests that DS microglia of the basal forebrain are activated. These results are consistent with reports of neuroinflammation, primarily from the cortex, as DS-AD progresses (89). Protein folding and protein maturation processes are downregulated in DS basal forebrain oligodendrocytes, suggesting that DS oligodendrocytes have impaired production of myelin components, possibly contributing to the reduced myelination of neurons in DS(56, 67, 68). Future work will need to validate these cell specific transcriptomic changes with biochemical assays and elucidate how the non-neuronal cells contribute to basal forebrain pathology and early degeneration of DS BFCNs.

### 4.6 Future Directions

Taken as a whole, our results reveal that metabolic dysfunction and oxidative stress are present in the DS basal forebrain by birth. Sustained metabolic dysregulation and exposure to oxidative stress from birth likely contributes to the susceptibility of BFCNs so early in the progression of DS-AD in individuals with DS. Elucidation of the shift in energy metabolism across the DS lifespan with human basal forebrain tissue or in stem cell models of BFCNs (90–95) will validate the transcriptomic changes. If these metabolic changes hold true, regulation of the OXPHOS pathway and ROS accumulation could provide targets for early therapeutic intervention in DS BFCNs prior to degeneration.

## 6. Acknowledgements

Human tissue was obtained from the NIH NeuroBioBank at the University of Maryland, Baltimore, MD. N.R.W. and A.B. would like to acknowledge the contributions of Bhattacharyya Lab members Dr. Matthew Russo, Grace Branger, Kamryn Witkowiak, Rachel Lichte, and Sahith Puthireddy for their preliminary stereological analysis of the basal forebrain that played a key role in the conceptualization of this project. We would like the acknowledge the feedback on this study provided by members of Dr. Su-Chun Zhang’s lab, members of Dr. Andre Sousa’s lab, Dr. Xinyu Zhao, Dr. Luigi Puglielli, Dr. Elizabeth Head, and Dr. Carissa Sirois. N.R.W. received training in bioinformatic analysis from the INCLUDE Data Science for Developing Scholars in Down Syndrome (DS3) which is supported by the National Institutes of Health INCLUDE Project under R25HD114950 from the Eunice Kennedy Shriver National Institute of Child Health and Human Development.

## 7. Conflicts of Interest

The authors have no conflicts of interest to disclose.

## 8. Funding Sources

Research reported in this publication was supported by the National Institutes of Health under F99AG086925 from the Office of the Director, National Institutes of Health and the National Institute on Aging awarded to N.R.W.; GM140935 Medical Scientist Training Program T32 and F30MH140382 Ruth L. Kirschstein Individual Predoctoral NRSA from the National Institutes of Health awarded to R.D.R.; R36AG072108 from the National Institute on Aging awarded to J.L.M; R01HD106197 from the Eunice Kennedy Shriver National Institute of Child Health and Human Development awarded to S-C.Z., D.F., A.M.M.S., and A.B; and P50HD105353 from the Eunice Kennedy Shriver National Institute of Child Health and Human Development awarded as a core grant to the Waisman Center. Additional funding came from the University of Wisconsin-Madison Sophomore Research Fellowship awarded to S.M.; BRFSG-2023-11 from the Brain Research Foundation awarded to A.M.M.S; and the William F. Vilas Trust Estate Award, Wisconsin Alzheimer’s Disease Research Center Research Education Component Award, and the Wisconsin Partnership Program New Investigator Award from the University of Wisconsin-Madison to A.B. The content of this study is solely the responsibility of the authors and does not necessarily represent the official views of the National Institutes of Health.

## 9. Consent Statement

Acquisition of the de-identified human tissue samples was approved by the Health Sciences Institutional Review Board at the University of Wisconsin-Madison (Protocol #2016-0979) and certified exempt from human subjects IRB oversight.

## Notes

### Competing Interest Statement

The authors have declared no competing interest.

### Summary of Updates

The results have been revised to reflect new analysis.

