## Supplemental Figures for "Single-nucleus analysis reveals oxidative stress in Down syndrome basal forebrain neurons at birth"

### Supplement 1

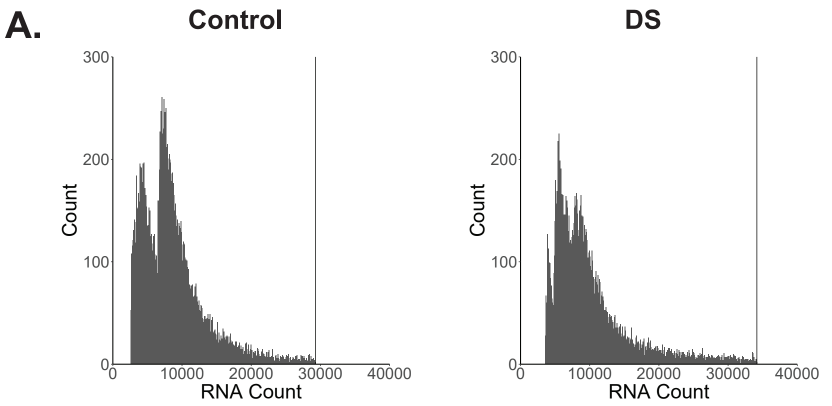


C.

| | Control | DS |
|---|---|---|
| Doublets | 1067 | 375 |
| Low Quality | 464 | 265 |
| **Total Cells Removed** | 1531 | 640 |

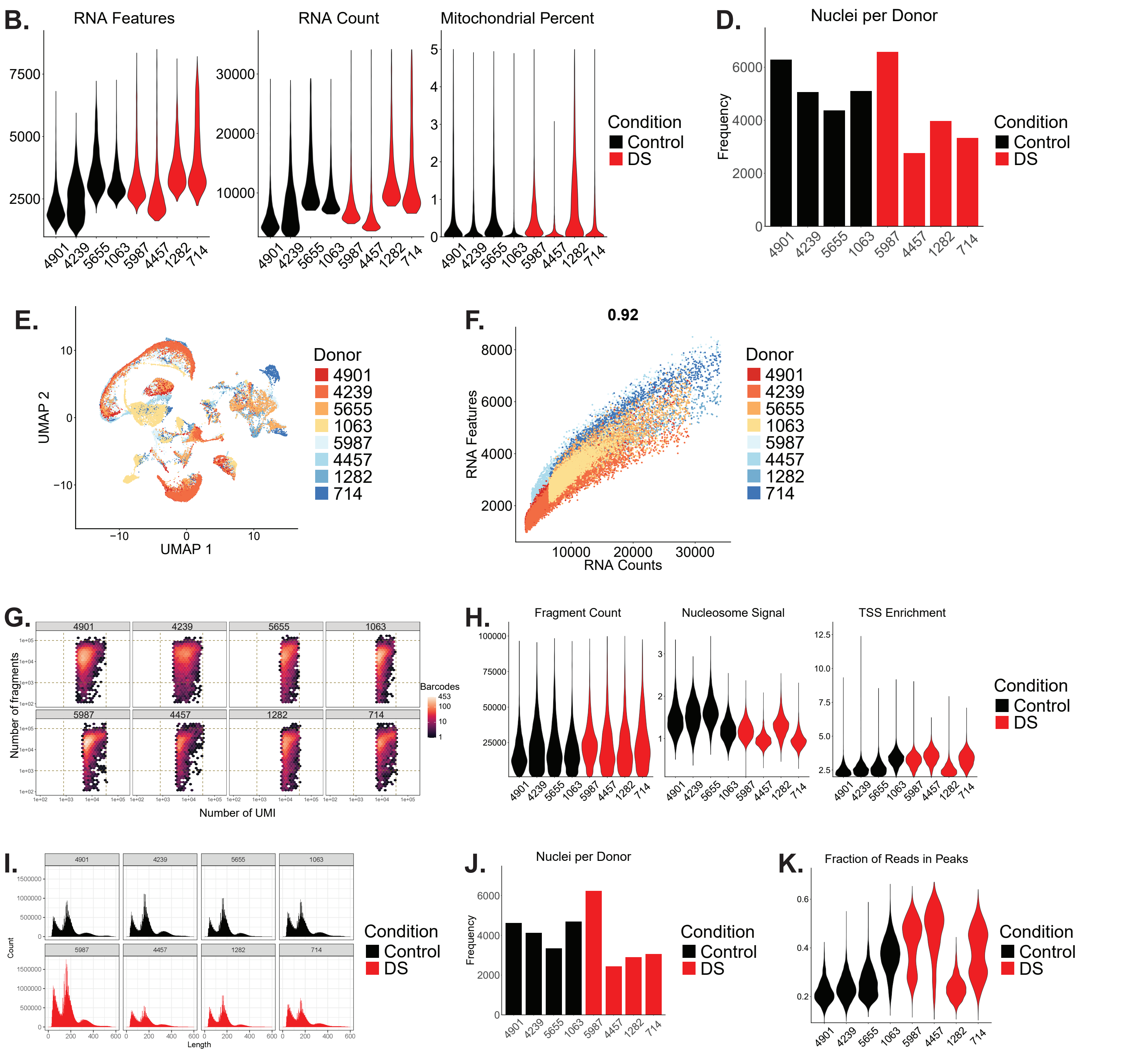

### Supplement 2

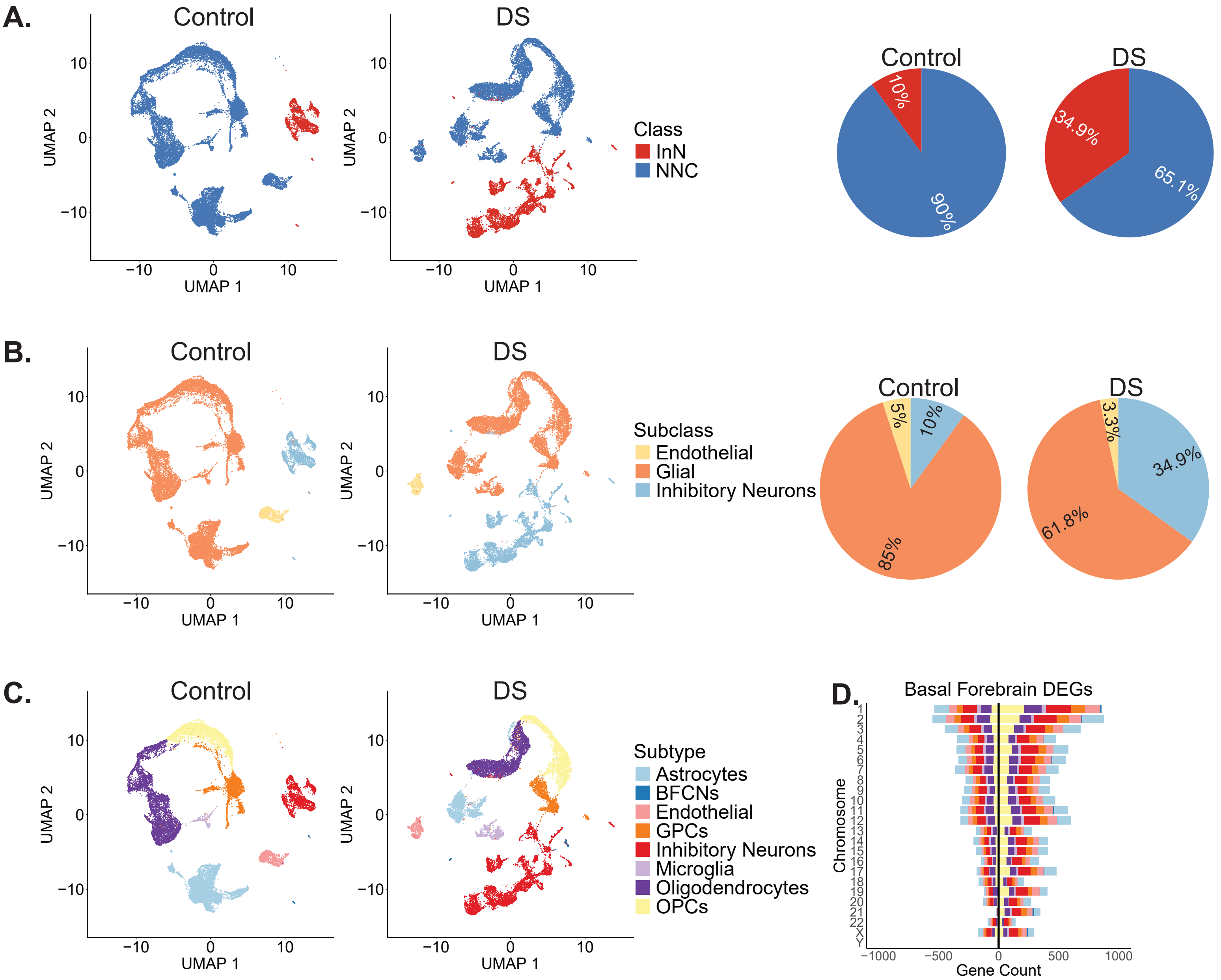

### Supplement 3

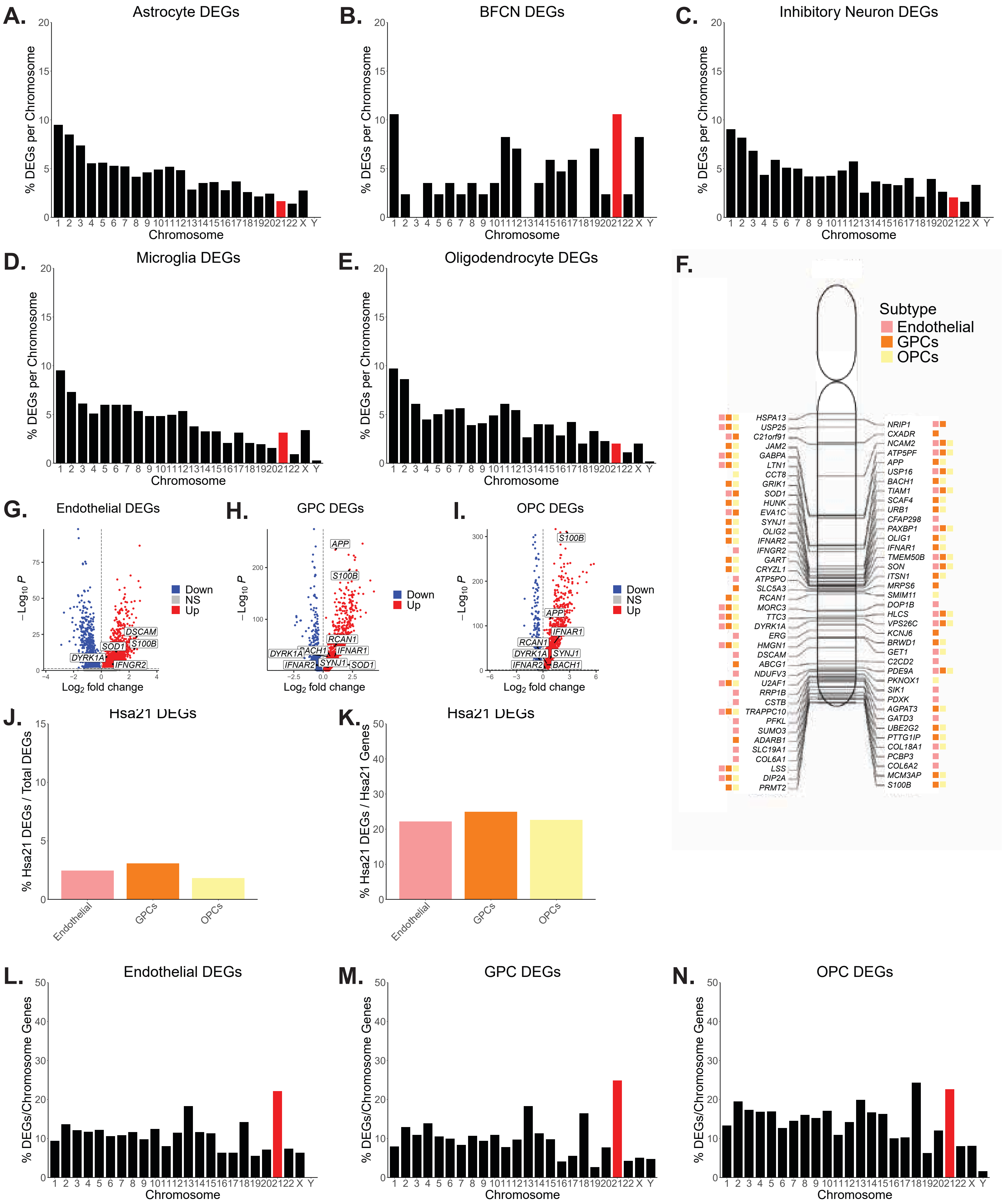

### Supplement 4

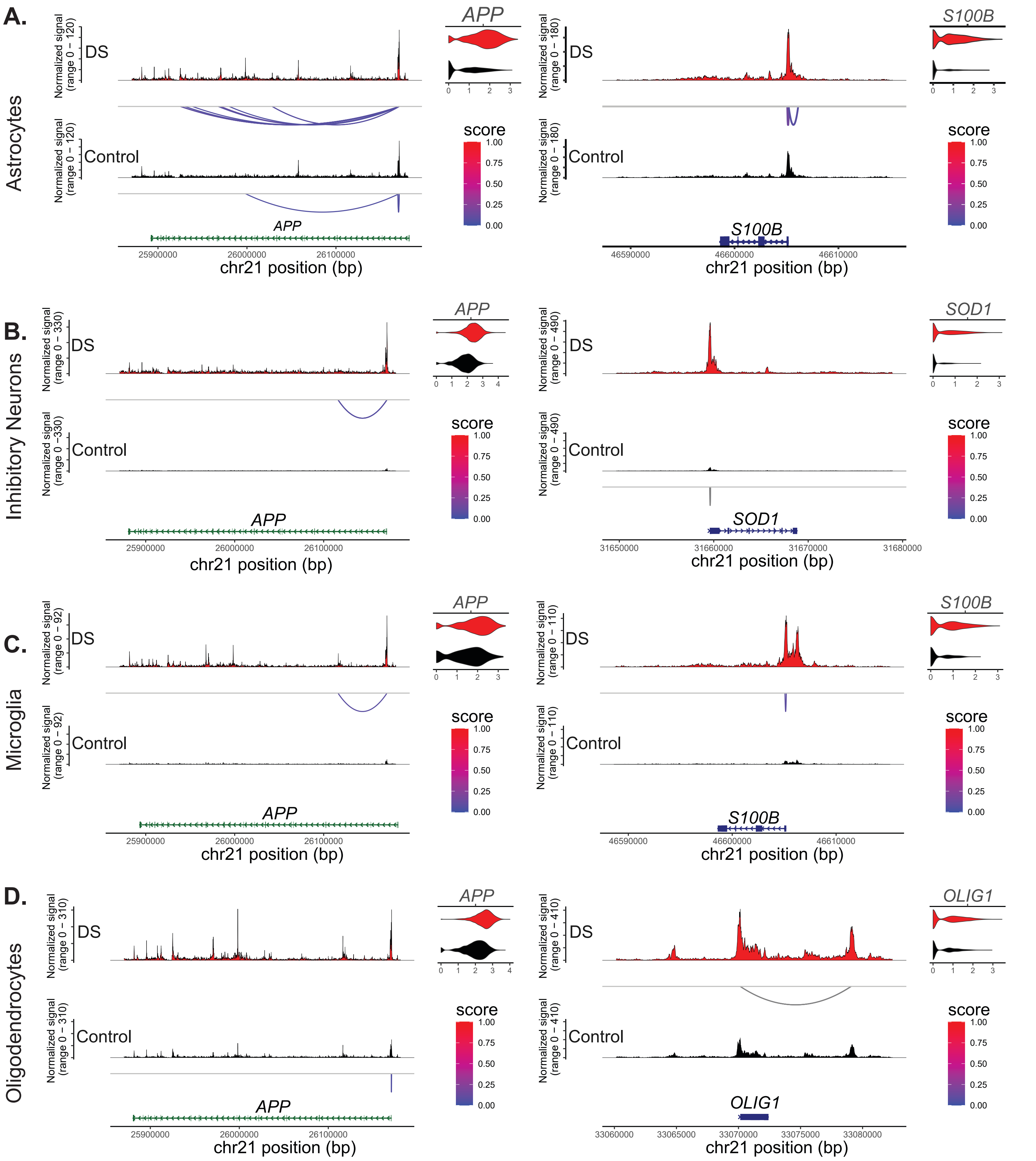

### Supplement 5

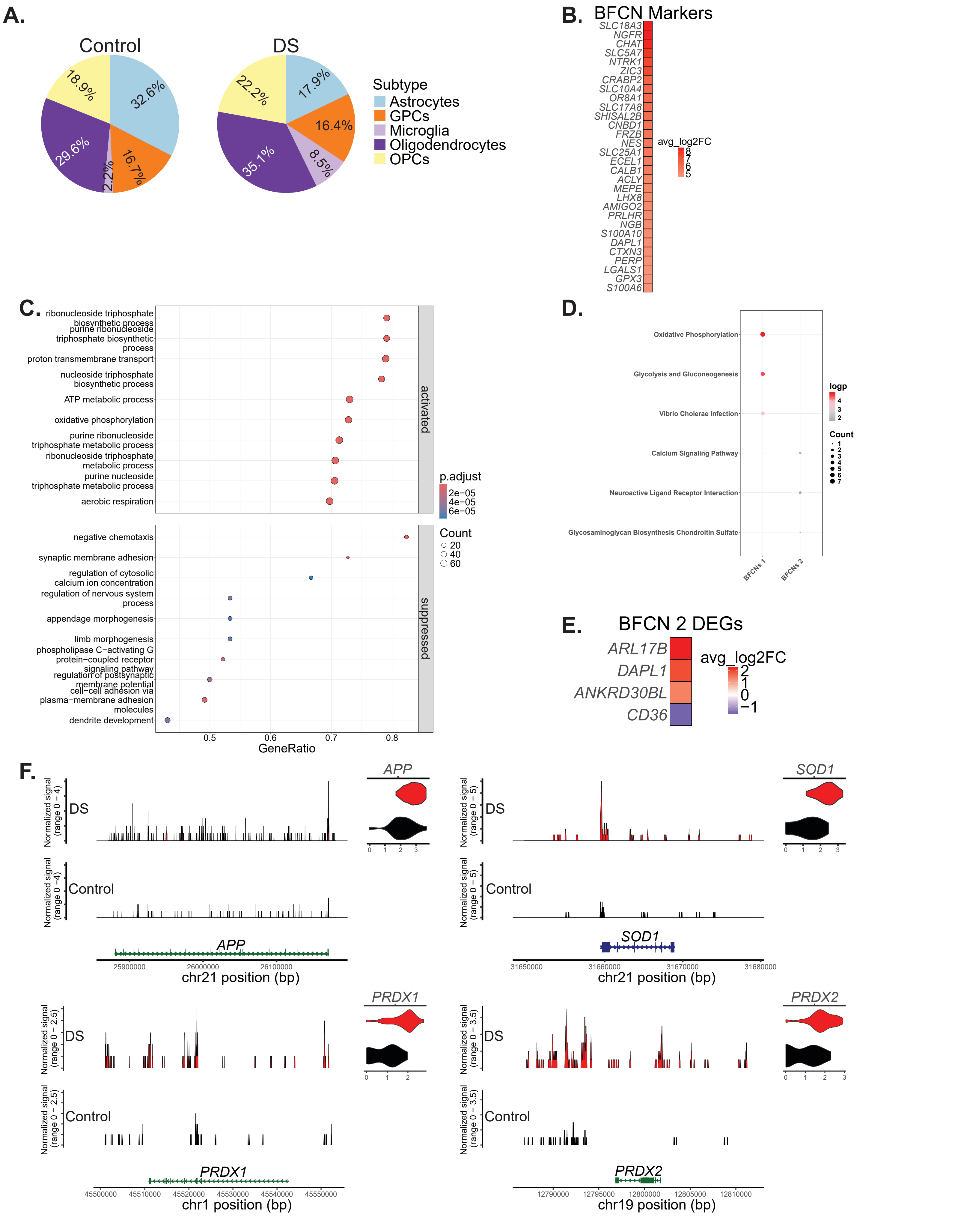
